# Human dorsal anterior cingulate neurons signal conflict by amplifying task-relevant information

**DOI:** 10.1101/2020.03.14.991745

**Authors:** R. Becket Ebitz, Elliot H. Smith, Guillermo Horga, Catherine A. Schevon, Mark J. Yates, Guy M. McKhann, Matthew M. Botvinick, Sameer A. Sheth, Benjamin Y. Hayden

## Abstract

Hemodynamic activity in dorsal anterior cingulate cortex (dACC) correlates with conflict, suggesting it contributes to conflict processing. This correlation could be explained by multiple neural processes that can be disambiguated by population firing rates patterns. We used *targeted dimensionality reduction* to characterize activity of populations of single dACC neurons as humans performed a task that manipulates two forms of conflict. Although conflict enhanced firing rates, this enhancement did not come from a discrete population of domain-general conflict-encoding neurons, nor from a distinct conflict-encoding response axis. Nor was it the epiphenomenal consequence of simultaneous coactivation of action plans. Instead, conflict amplified the task-relevant information encoded across the neuronal population. Effects of conflict were weaker and more heterogeneous in the dorsolateral prefrontal cortex (dlPFC), suggesting that dACC’s role in conflict processing may be somewhat specialized. Overall, these results support the theory that conflict biases competition between sensorimotor transformation processes occurring in dACC.

## INTRODUCTION

When faced with conflicting demands for attention or action, we can marshal cognitive resources to maintain effective performance despite this conflict (Shenhav et al., 2013; Botvinick and Braver, 2015; Botvinick and Cohen, 2014; Kerns et al., 2004; Shenhav et al., 2017). The ability to respond adaptively to conflict is a hallmark of higher cognition, one that allows us to devote the appropriate level of cognitive resources to make good decisions. However, the way in which the brain detects and resolves conflict is a poorly understood aspect of higher-level cognition.

Conflict processing is often associated with the dorsal anterior cingulate cortex (dACC), a region in which conflict alters or increases brain activity (Botvinick et al., 1999; Botvinick et al., 2001 Shenhav et al., 2016). There has been a long-running debate about what effect, if any, conflict has on neuronal computations (Cole et al., 2009). This debate is driven in part by prominent failures to observe conflict correlates at the single unit level, (Amiez et al., 2005; Amiez et al., 2006; Blanchard and Hayden, 2014; Cai and Padoa-Schioppa, 2012; Ito et al., 2003; Nakamura et al., 2005). However, more recent studies have shown single neuron correlates of conflict in non-human animals (Ebitz and Platt, 2015; Bryden et al., 2018; Michelet et al., 2015). Most importantly, studies in human dACC – which lack translational uncertainties associated with model species – provide unambiguous correlates of conflict in both single units and in local field potentials (Sheth et al., 2012; Smith et al., 2019). These results confirm that conflict has direct and measurable neuronal effects but leave unresolved the computations underlying these effects. Here, compare three possibilities, each consistent with recent discoveries, in a dataset collected in humans performing a conflict task. (Note that these three hypotheses are not necessarily mutually exclusive.)

The ***explicit hypothesis*** proposes that dACC neurons signal conflict abstractly, in the sense that conflict-related modulations serve the purpose of transmitting information about the presence of conflict – in general – to downstream conflict resolution structures, which implement its resolution. These downstream structures likely include the dorsolateral prefrontal cortex (dlPFC, Johnston et al., 2007; Ma et al., 2019; Smith et al., 2019; MacDonald et al., 2000; Shenhav et al., 2013). In this view, dACC contains either a dedicated, discrete set of neurons specialized for encoding conflict or else its neurons have a distinct population coding *axis* (sometimes referred to as a *dimension*, i.e. some linear combination of neuronal responses) that encodes conflict via small, distributed changes across a large number of neurons.

The ***epiphenomenal hypothesis*** proposes that conflict correlates are the epiphenomenal consequence of the co-activation of neurons that are tuned for different actions (Nakamura et al., 2005) or response predictions (Alexander and Brown, 2011). Epiphenomenal, here, means that correlates of conflict are not driven by computations related to conflict *per se*, but nonetheless covary with it. This hypothesis was first motivated by the prominent failures to find unit correlates of conflict in a pioneering study of macaque supplementary eye field (SEF), a structure adjacent to dACC (Nakamura et al., 2005). Nakamura and colleagues found that neuronal correlates of conflict in SEF can be explained by co-activation of sets of neurons selective for basic task variables. It is possible that the same ideas may apply to dACC, as goes this hypothesis.

The ***amplification hypothesis*** proposes that conflict does affect dACC neurons, but does so by amplifying task-relevant information encoded in dACC neurons. This view is motivated by two observations. First, recent work has amply demonstrated that neurons in dACC are robustly tuned for a variety of sensory and motor variables (Heilbronner and Hayden, 2016), so the region has all the requisite signals to directly participate in sensorimotor transformations. Second, the ultimate function of conflict processing is not to detect conflict, but to resolve it. One natural way to do so is to amplify task-relevant sensorimotor information at the expense of irrelevant information (Shenhav et al., 2013; Egner and Hirsh, 2005; Botvinick and Cohen, 2014).

To arbitrate between these three hypotheses, we examined a large dataset of single neurons recorded in human dACC and, for comparison, a complementary dataset recorded in human dlPFC. Participants performed the multi-source interference (MSIT) task that independently manipulates two forms of conflict, Simon (motor) and Eriksen (perceptual). We find that both forms of conflict modulate responses of single dACC neurons and both tend to increase average firing rates. However, the epiphenomenal hypothesis could not account for neural responses in dACC, and our dimensionality reduction results were more consistent with the amplification hypothesis than with the explicit hypothesis. Our results indicate that conflict robustly enhances the strength of coding of task-relevant sensorimotor information by shifting patterns of population activity along coding dimensions that correspond to the identity of the correct response. This pattern is predicted by the conflict amplification hypothesis. Neurons in dlPFC respond considerably more weakly and heterogeneously to conflict, suggesting that dACC may have a relatively specialized role in conflict processing.

## RESULTS

We examined neuronal responses collected from 16 human subjects (dACC: n=7 patients, dlPFC: n=9 patients, see **Methods**) performing the multi-source interference task (MSIT; **Figure 1A-B**). This task and its close variants have been widely used to study conflict in humans in studies using both mass action measures and intracranial electrophysiology (Sheth et al., 2012; Smith et al., 2019; Widge et al., 2019a). These data were recorded in human dACC and dlPFC (**Figure 1C**). Some of these data come from a set used in a previous publication that focused on local field potentials, which are not relevant to our hypotheses and are not considered here (Smith et al., 2019). The data we study here do not overlap with those used in Sheth et al. (2012), although the tasks are identical.

**Figure 1.**
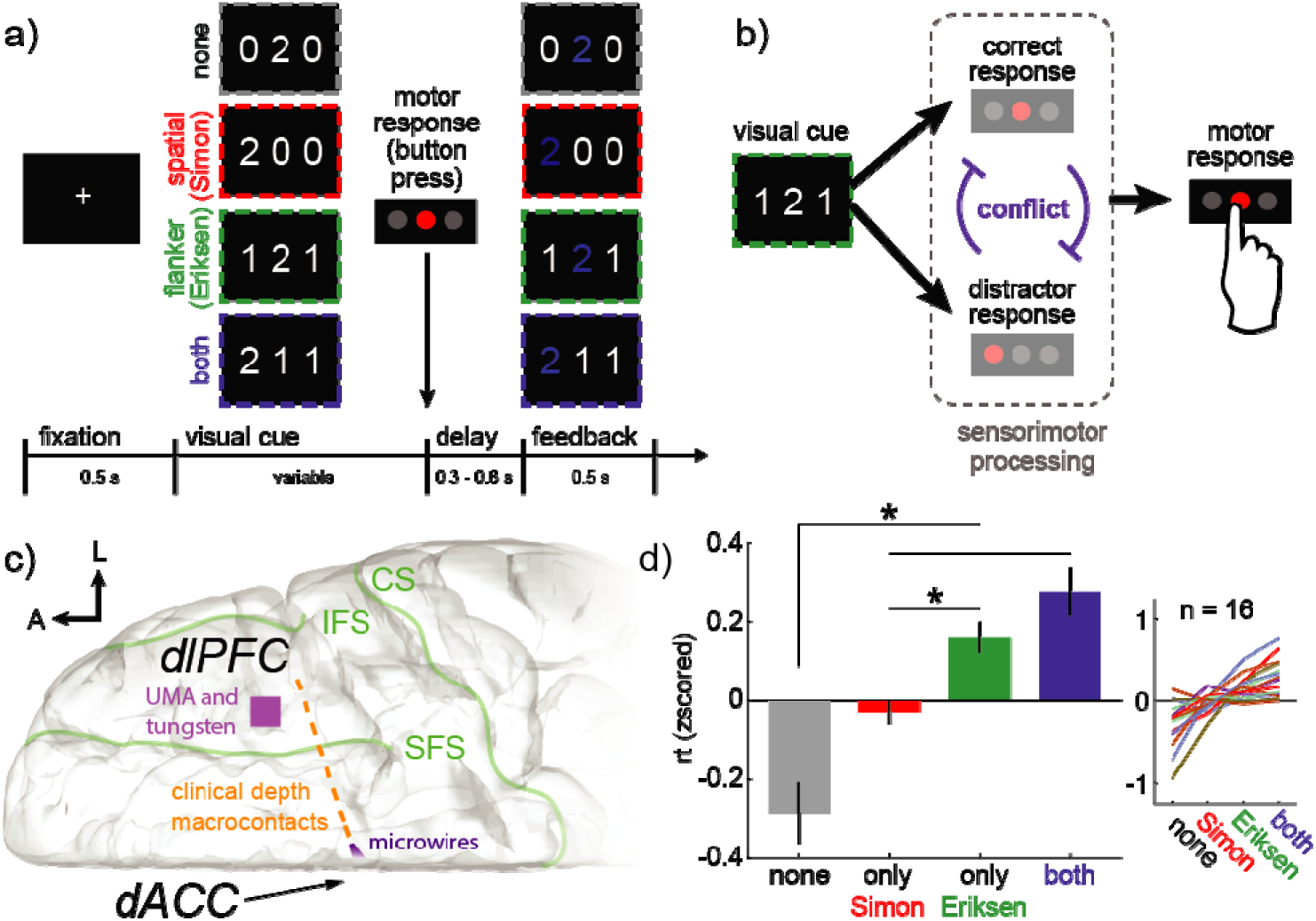
MSIT task and anatomy: **A.** Structure of the multi-source interference task (MSIT). The subject sees a visual cue consisting of 3 numbers and has to identify the unique number with a button push. The “correct response” is the left button if the target is 1, middle if 2, right if 3. Four example cues are shown here, and in each case, the target is “2” and the middle button is the correct response. This is most obvious for the first cue (“none”), where there is no conflicting information. In the other three examples, conflicting information makes the task more difficult. First, incongruence between the location of the target number in the 3-digit sequence and location of the correct button in the 3-button pad produces spatial (Simon) conflict (orange). Second, the distracting presence of numbers that are valid button choices (“1”, “2”, “3”) produces flanker (Eriksen) conflict (green). Trials can also simultaneously have both types (blue). **B.** The visual cues are associated with one or more sensorimotor responses. Every cue has a **correct response**, meaning the button press that corresponds to the unique target. Cues can also have one or more **distractor responses**, meaning the button press that corresponds to task-irrelevant spatial information (Simon) or flanking distractors (Eriksen). If and only if the correct response and distractor response do not match, then the cue causes **conflict** because only one button response can ultimately be chosen. **C.** Diagram of the intracranial implant including a stereotactically placed intra-cerebral depth electrode with macroelectrodes (blue squares) along the shaft from dlPFC to dACC and microwire electrodes (orange star) in dACC. A, anterior; L, lateral; CS, central sulcus; SFS, superior frontal sulcus; IFS, inferior frontal sulcus. The UMA and tungsten microelectrode recoding locations are schematized as a purple square on the surface of dlPFC. **D.** The average (mean) response times across subjects in each of the four task conditions and (right) the mean response times within each subject. Bars = standard error across subjects.

The MSIT independently manipulates two forms of conflict, either with flanking distractors (Ericksen conflict) or by using the discrepancy between the position of the task-relevant cue and the correct button press (Simon conflict). Response time was slower when any form of conflict was present (**Figure 1D**; mean z-scored response times: no conflict =-0.29 ± 0.07 STE across subjects; any conflict = 0.11 ± 0.03 STE; mean within-subject difference = 0.39 ± 0.1 STE; sig. difference, p < 0.002, t(2,14) = 4.02, paired t-test). The effects of Simon and Eriksen conflict appeared to be additive; greatest response time slowing occurred when both were present (mean response time for only Simon conflict trials =-0.03 ± 0.03 STE across subjects; only Eriksen conflict trials = 0.16 ± 0.04 STE; trials where both forms of conflict were present = 0.28 ± 0.06 STE). Further, response time was consistently slower during Eriksen conflict compared to Simon (mean within-subject difference = 0.19 ± 0.05 STE; paired t-test: p < 0.003, t(2,14) = 3.72), suggesting that Ericksen flankers were slightly more effective at driving conflict in this task.

### Encoding of conflict in single neurons in dACC

We recorded from 145 dACC neurons from 6 human patients. Because our previous investigations show that neural responses can be relatively long-lasting in dACC (Hayden et al., 2011), we chose a full-trial epoch analysis approach (specifically, a 3 second epoch starting at trial onset, roughly the duration of the trial). Note that we chose this analysis epoch before beginning data analysis. Example cells showing changes in firing rate associated with conflict are shown in **Figure 2A** and **Figure 2B**.

**Figure 2.**
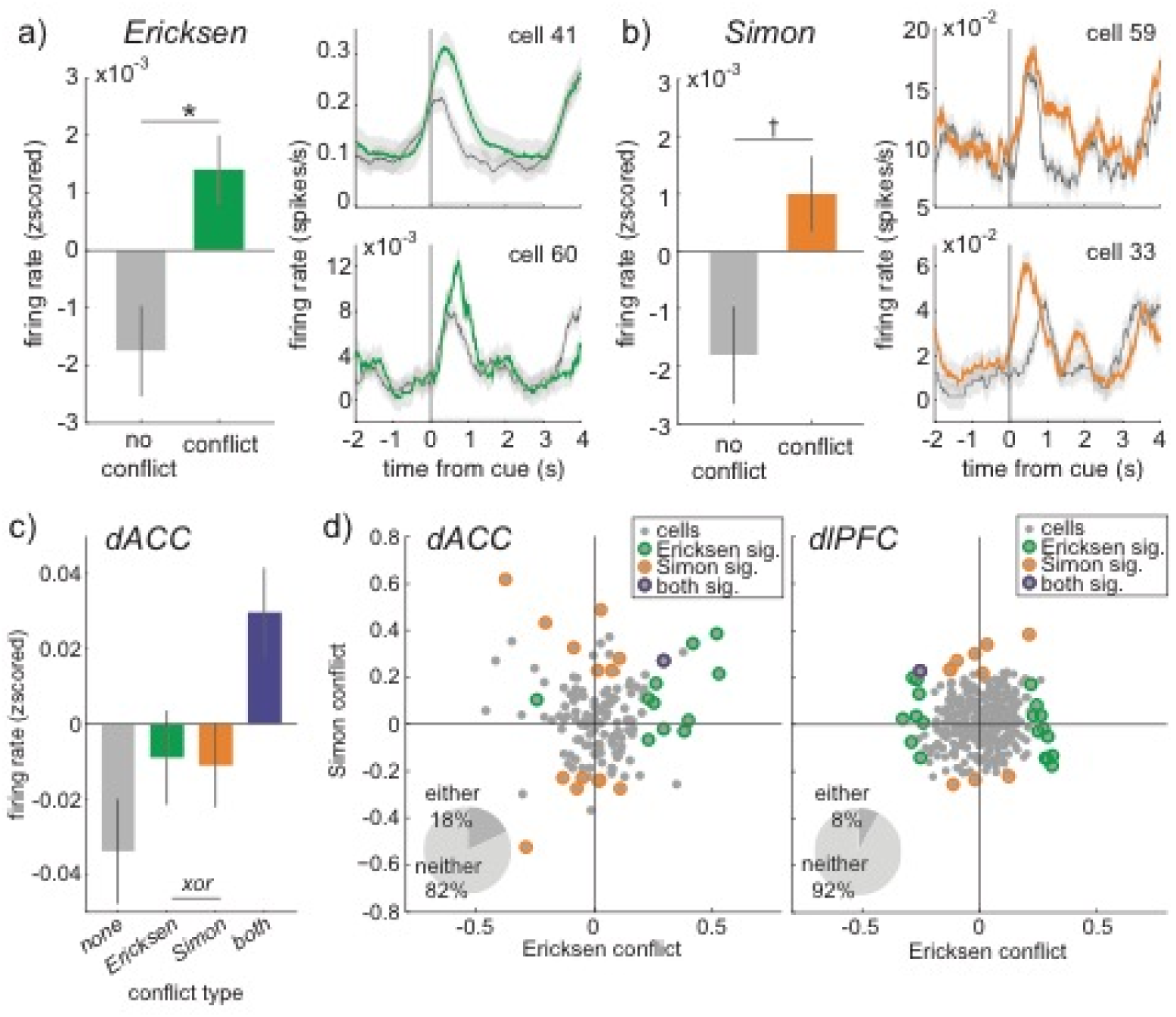
Additive effects of conflict at the population, but different conflict effects in single neurons. A) Left, Average firing rate across all neurons recorded in dACC during Ericksen conflict. Bars = STE, * p < 0.05, † p < 0.1. Right, Two example cells on no-conflict (gray) and Ericksen conflict trials (red). Ribbons = STE. B) Same as A, for Simon conflict (green). C) Additive effects of each type of conflict. D) Distribution of Simon and Ericksen conflict effects within single neurons in dACC (left) and dlPFC (right). Circled neurons respond significantly (p < 0.05) to the highlighted form of conflict (red = Ericksen; green = Simon; blue = both).

Across the population of dACC neurons, activity was higher on Ericksen conflict trials than on no conflict trials (**Figure 2A**; t-test for all neurons on all trials, p < 0.03; mean increase = 0.022 z-scored spikes/s ± 0.01 STE). A small number of individual neurons also had different activity levels on Eriksen conflict and no conflict trials (8.2%, n=12/145 neurons, within-cell t-test). This proportion is slightly greater than chance (p < 0.04, one-sided binomial test). In all but one of these neurons, conflict increased firing (significant positive bias, p < 0.0005, one-sided, binomial test; mean increase in these cells = 0.30 z-scored spikes/s ± 0.06 STE).

Simon conflict was also associated with an increase in activity across the population of dACC neurons, although this increase was not statistically significant (**Figure 2B**; p < 0.06, mean increase = 0.0007, z-scored spikes/s ± 0.0004 STE). Overall, 10.3% (n=15/145) neurons had significantly different firing rates between Simon and no-conflict trials. This proportion is greater than chance (p < 0.003, binomial test). However, the sign of conflict encoding in these cells was nearly even (8/15 showed increasing activity; mean increase in these cells = 0.06 z-scored spikes/s ± 0.09 STE). This result indicates that, while dACC neurons do encode Simon conflict, the effect is not strongly (if at all) directional, unlike the positive bias we observed for Eriksen conflict (see above).

The largest increase in activity occurred on trials that induced both Simon and Ericksen conflict. Model comparison revealed that the effects of Simon and Ericksen conflict were essentially equivalent and, again, additive across the population of neurons (**Figure 2C**; **Table S1**). An additive model was a better fit to the data than other, more flexible models (all BIC weights < 0.02; sig. additive term: β_1_ = 0.033, p < 0.003; see **Methods**). Thus, though Simon conflict was perhaps more weakly encoded in dACC than Ericksen conflict, regardless of its source, conflict mostly increased the activity of dACC neurons and the effects of Simon and Ericksen conflict were additive.

We recorded responses of 378 neurons in dlPFC from 9 patients. In contrast to dACC, we observed little modulation by either form of conflict in dlPFC. Across dlPFC neurons, activity was not higher during Ericksen conflict trials, compared to no-conflict trials (p > 0.5, paired t-test; mean increase < 0.0001 z-scored spikes/s ± 0.0002 STE). Average firing rate was higher during Simon conflict, but the effect size was very small (p < 0.005; mean increase = 0.0005 z-scored spikes/s ± 0.0002 STE). Both effect sizes were significantly smaller than the corresponding effects measured in dACC (p<0.001, t-test). The number of individual neurons that showed individual conflict-related modulation did not significantly exceed the expected false positive rate of 5% (Ericksen conflict: 5.3%, n=20/378; Simon conflict: 2.9% n=11/378 neurons). Fewer neurons had any tuning for either form of conflict in dlPFC, compared to dACC (p < 0.05; compare dlPFC: 7.9%, n=30/378 cells; dACC: 17.9%, 26/145 cells; two-sample proportion test, pooled variance). Thus, while conflict responses in dACC were weak, they were larger in dACC than in dlPFC, and responses were more consistently positive in dACC than in dlPFC.

### Simon and Ericksen conflict tend to affect distinct pools of neurons

We next considered whether neurons in dACC carry an abstract conflict signal, that is, one that indicates the presence of conflict, regardless of its source. If dACC detects conflict, then individual dACC neurons that are sensitive to Ericksen conflict should also be sensitive to Simon conflict. Our data do not support this idea. Simon and Ericksen conflict had unrelated effects on individual neurons. That is, we observed no significant correlation between the modulation indices for Simon and Ericksen conflict (r = 0.05, p > 0.5; **Figure 2D**). Furthermore, the population of cells whose responses were significantly affected by Eriksen conflict was almost entirely non-overlapping with the population significantly affected by Simon conflict (specifically, only one cell was significantly modulated by both). The proportion of co-activated dACC neurons was not substantively different from what we observed in dlPFC (n = 1/378 cells significant for both forms of conflict in dlPFC; no difference in proportions, p > 0.4, two-sample proportion test with pooled variance). The correlation between Simon and Ericksen conflict responses in dlPFC neurons (r = 0.06, p > 0.2) also closely matched the values found in dACC. Thus, we found no evidence that dACC neurons uniquely carried some abstract conflict signal. In other words, our evidence does not support the idea that dACC carries conflict-related information that is non-specific to the type of conflict.

### Conflict coding in neurons is not epiphenomenonal

We next considered whether conflict encoding was an epiphenomenal consequence of co-activating pools of neurons tuned for different stimuli and/or action plans. This idea was originally proposed by Nakamura et al. (2005). For simplicity, we use the term “response tuning” to indicate selectivity for the sensorimotor responses that were required for the task (“correct responses”), agnostic to whether this tuning was at the level of cue processing, generating the button box response, or the transformation from cue to response. We use the term “distractor response”, to refer to the conflicting sensorimotor response indicated by the conflicting cues.

Nakamura’s epiphenomenal hypothesis predicts that there are separate pools of neurons corresponding to the two conflicting actions, and that conflict increases activity because it uniquely activates both pools. We used ANOVA to jointly estimate the effects of the correct responses, distractor responses, and the conflict between the two on the firing rates of dACC neurons (**Figure 3A**; see Methods). We found that responses of a significant proportion of neurons were selective for the correct response (13.1% ± 2.8% STE, n = 19/145 neurons, this proportion is greater than chance, 5%, p < 0.0001, one-sided binomial test). However, neurons did not encode the distractor response (because we considered tuning for either Ericksen or Simon distractors, the chance level false positive rate was 9.75%; percent significant cells 9.7% ± 2.5% STE, 14/145 neurons, p > 0.4, one-sided binomial test against chance). Despite the fact that few neurons encoded the distractor response, a significant proportion of neurons did still signal either Ericksen or Simon conflict (16.6% ± 2.5% STE, n = 24/145 neurons, greater than chance at 9.75%, p < 0.004). Thus, conflict signals occurred *more* frequently in single neurons than we would expect from the epiphenomenal conflict view, where conflict could only emerge in neurons tuned for both correct and distractor responses.

**Figure 3.**
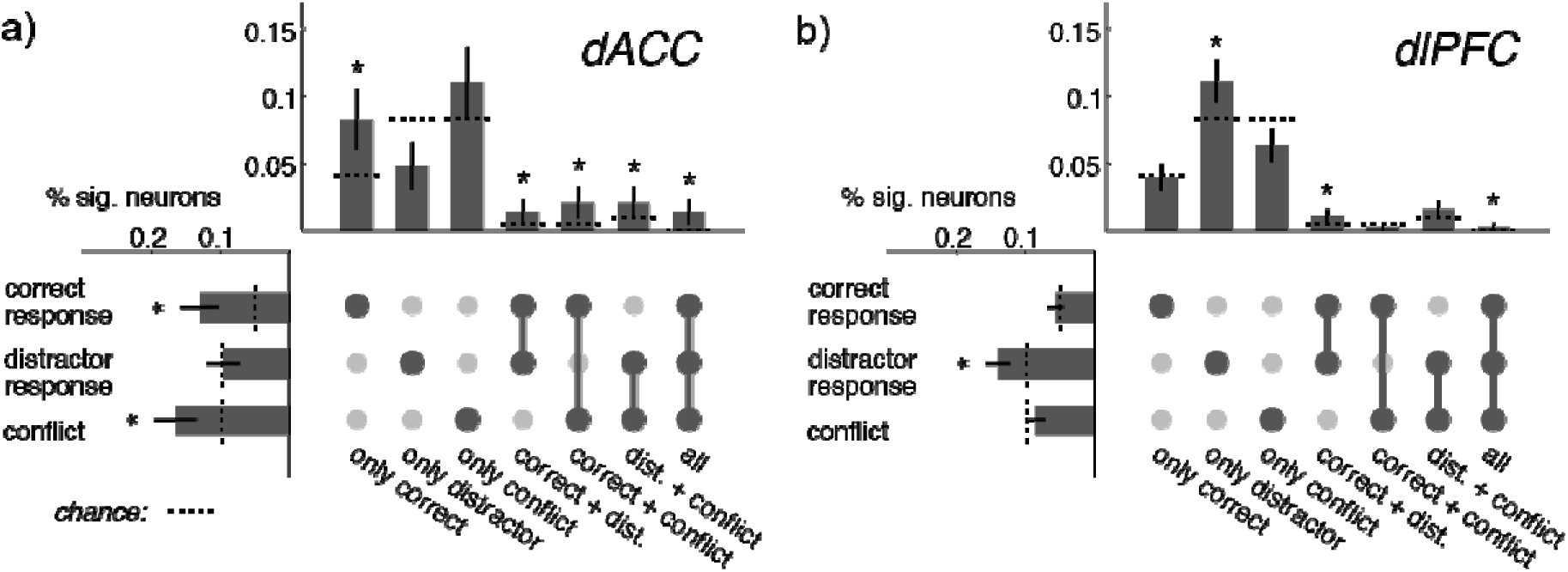
Relationships between task, distractor, and conflict tuning in dACC neurons. A) Percent neurons significantly tuned for task, distractor, conflict (left) and combinations of these variables (top) in dACC. Dotted lines reflect expected false positive rates for each condition. Bars = STE, * p < 0.05 greater than false positive rate. B) Same as A for dlPFC.

More critically, even in correct response-selective neurons, the preferred correct response rarely matched their preferred Simon/Ericksen distractor response (only 5.3% of cells matched, ± 1.9% STE, one sided binomial test against chance at 11%, p = 1). The epiphenomenon hypothesis would predict 100% match. Moreover, while a very small proportion of cells were response tuned for both correct responses and distractor responses; 2.8% ± 0.1% STE, 4/145), the majority of conflict-modulated dACC neurons came from a different set (91.7% ± 0.2% STE, n = 22/24). In fact, the majority of conflict-sensitive dACC neurons were not selective for either correct response or distractor responses (66.7% ± 0.3% STE, n = 16/24) – a result that is in direct opposition to the idea that these signals are an emergent consequence of response tuned cells. Thus, not only were response tuned neurons not responsible for the majority of conflict signals in dACC, but the data did not support even the basic premise that there were generic response tuned neurons in dACC.

In dlPFC (**Figure 3B**), conversely, neurons were selective for distractor responses (14.0% ± 1.8% STE, 53/378 neurons, greater than chance at 9.75%, p < 0.004). Like dACC, few neurons were selective for the combination of correct responses and distractor responses (1.3%, 5/378 neurons, sig. greater than chance at 0.5%, p < 0.02), but in dlPFC these responses matched. Neurons that were tuned for a specific correct response were often tuned to prefer the *same* Simon/Ericksen distractor response (19% of cells matched, ± 2.0% STE, one sided binomial test against chance at 11%, p < 0.0001). Thus, we did see some evidence of generic response tuning in dlPFC, but not in dACC. However, unlike dACC, there was not substantial selectivity for conflict in dlPFC (8.5% ± 1.4%, compare to chance at 9.75%, n = 32/378; correct response = 5.6% ± 1.2%, compare to chance at 5%, n = 21/378). Ultimately, although the tuning properties of dlPFC neurons were more likely to match the premises of the epiphenomenal hypothesis than dACC neurons, dACC neurons, not dlPFC neurons, were more likely to signal conflict.

### Population analyses can cleanly disambiguate our hypotheses

At the population level, the three hypotheses make different predictions for neural activity. The **explicit hypothesis** predicts that there should be either a set of conflict-selective neurons or there should be a conflict-selective *axis* in the population. A population axis is, by our definition here, some linear combination of neuronal firing rates that tracks the presence or absence of conflict, but is distinct from any other parameter the population may encode. Note that the former, stronger prediction (a subset of conflict-selective cells) would also satisfy the latter, weaker prediction (a conflict-encoding axis), so we focus on the latter prediction to maximize the chances of validating this model. In the **epiphenomenon hypothesis**, when correct response and distractor responses match (i.e., when there is no conflict), both inputs activate the same set of neurons (**Figure 4A**, left). When they are in conflict, separate sets of neurons are activated (**Figure 4A**, right, Nakamura et al., 2005). At the population level, then, the **epiphenomenon hypothesis** predicts that conflict should *decrease* the amount of information about the correct response and shift neuronal population activity down along the axis in firing rate space that encodes this response (**Figure 4B**). Note that net population activity will only increase if conflict increases activity in the distractor response neurons more than it decreases activity in the correct response neurons (Nakamura et al., 2005). As a result, in the **epiphenomenal hypothesis**, as in the **explicit hypothesis**, there will be a population axis that selectively encodes conflict, corresponding to the summed activity of all the neurons. However, in the explicit case, this shift will only be in the direction of a unified conflict detection axis, whereas in the epiphenomenal view, it will largely, but not exclusively be along the coding dimensions in firing rate space that discriminate one response from another (**Figure 4C and D**). The **amplification hypothesis**, conversely, does not predict a unified conflict detection axis in the population. Instead, it makes a prediction that is exactly contrary to the epiphenomenal view: that conflict should shift population activity along task-variable coding dimensions, but in the opposite direction. That is, conflict is predicted to amplify task-relevant neural responses (**Figure 4E**).

**Figure 4.**
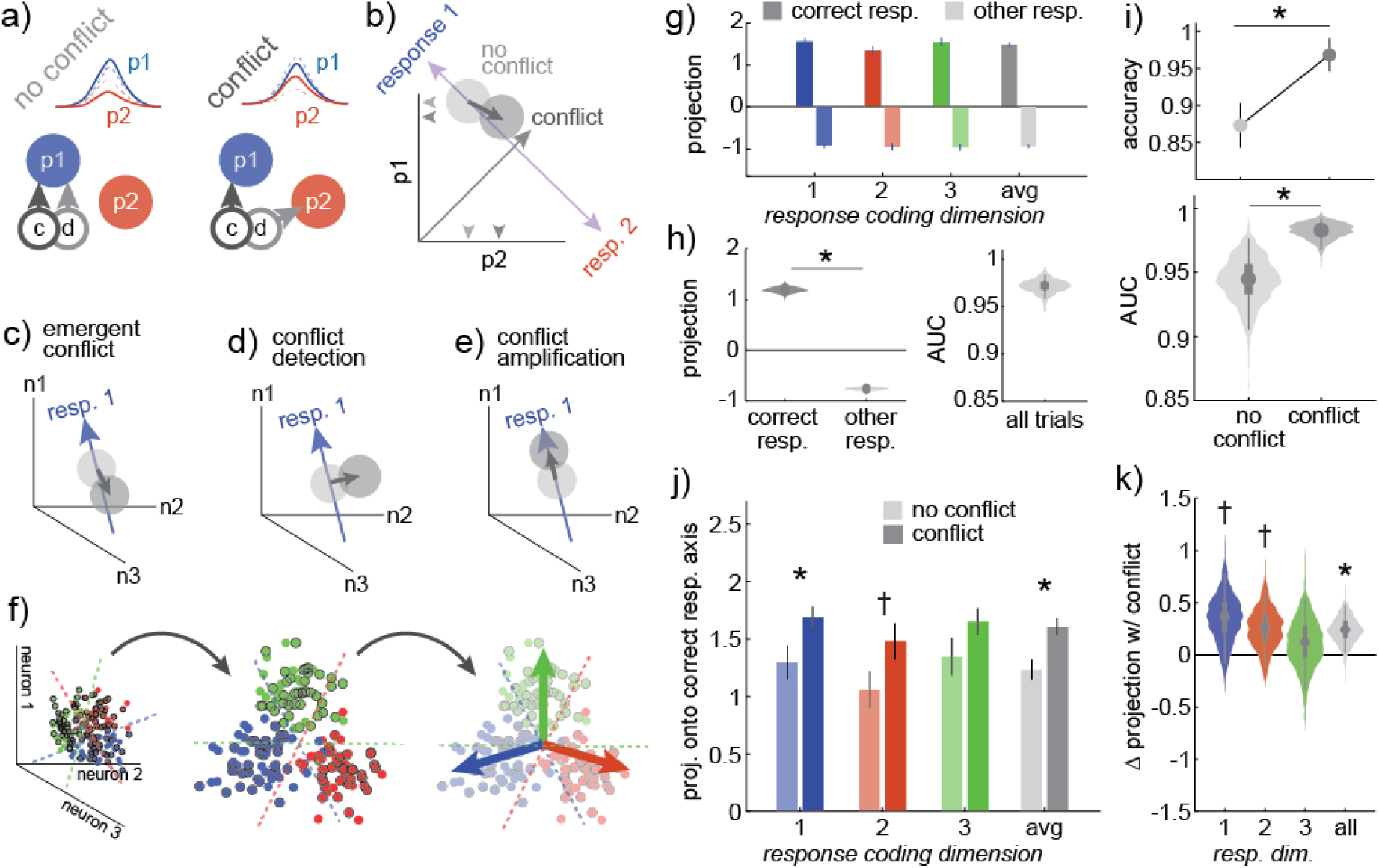
Population-level analyses suggest that dACC conflict signals amplify task representations. A) Cartoon of the epiphenomenal conflict hypothesis, where separate pools of neurons are tuned for response 1 (p1, blue) and 2 (p2, red). When the correct response is response 1 and there is no conflict, correct response (c) and distractor response (d) information both activate p1. When there is conflict, distractor information increases p2 activity at the expense of p1. If conflict increases p2 activity more than it decreases p1, total neural activity will be higher during conflict. B) A population view of the epiphenomenal conflict hypothesis. Here, p1 and p2 activity form the axes of a firing rate space, in which trials are distributed (shaded circles). In this firing rate space, there is a coding dimension that differentiates neural activity for correct response 1 (correct response = 1, regardless of conflict) from neural activity for the other responses, here response 2. This coding dimension is p1 – p2 here. In the epiphenomenal hypothesis, conflict decreases p1 activity and increases p2, which would largely shift response 1 activity down along the response coding dimension that differentiates response 1 from other responses. A conflict signal is epiphenomenal if activity also moves above this manifold, along an orthogonal, conflict-detecting axis (here, p1 + p2). C) The epiphenomenal hypothesis predicts that conflict should mostly, though not exclusively, shift activity down response coding dimensions, because it decreases the encoding of the correct response in favor of the distractor response. D) The explicit hypothesis predicts that conflict should largely shift activity along some conflict-detecting dimension that is orthogonal to response coding. E) The amplification hypothesis predicts that conflict should amplify the representation of response information—shifting activity up the response coding dimensions. F) Targeted dimensionality reduction to find response coding dimensions in the data. We find the separating hyperplanes that discriminate each response from the other two (left), project individual conflict (black circle) and no-conflict (no outline) trials into the subspace defined by these separating hyperplanes (middle), and measure projections onto the resulting response coding dimensions (right; pale arrows). G) Projections of one representative pseudopopulation onto the coding dimension that corresponds to the correct response that subjects executed on the trial (saturated), or to the other two responses (light). H) Distribution of mean projections onto the correct response and the other responses across pseudopopulations (left) and the distribution of AUC values for discriminating the correct response from the other responses based on these projections (right). I) Top: Task classification accuracy from coding dimension projections for conflict and no-conflict trials. One representative pseudopopulation. Bottom: Average AUC values for conflict and no conflict trials across pseudopopulations. J) Projections of conflict and no-conflict trials onto the correct response coding dimensions. H) Difference in correct response coding dimension projections between conflict and no conflict trials across pseudopopulations. Bars ± SEM and box plots illustrate the median, 50% and 95% CI. * = p < 0.05, † = p < 0.1.

### Conflict amplifies encoding of correct response information dACC

To arbitrate between these hypotheses, we must determine where trials fall along the coding dimensions for each correct response, where the “coding dimensions” are the combinations of neuronal firing rates that best predict the likelihood that the subjects are performing one of the three motor responses. We did this by combining the responses of neurons recorded separately into pseudopopulations (Churchland et al., 2012; Machens et al., 2010; Mante et al., 2013; Meyers et al., 2008; Ebitz et al., 2019) and then using a form of targeted dimensionality reduction to identify the coding dimensions in the population activity (Ebitz et al., 2018; Ebitz et al., 2019; Peixoto et al., 2019; Cunningham and Yu, 2014). Briefly, we used multiple logistic regression to identify the linear combinations of neuronal firing rates that encoded specific correct responses, then projected the activity from individual trials onto each coding dimension, that is, into the subspace defined by the coding dimensions corresponding to the three different responses (**Figure 4F**; see **Methods**).

When population activity was projected into task-coding space, it was easy to predict the current correct response from neural activity (**Figure 4G**-**H**; across 1000 bootstrapped populations: mean projection onto correct response coding dimension = 1.19, 95% CI = [1.08,1.30]; mean projection onto other response dimensions = −0.76, 95% CI = [−0.82, -0.70]; mean AUC = 0.97, 95% CI = [0.96,0.98]). However, classification accuracy was even higher for trials with Ericksen conflict than it was for trials without Ericksen conflict (**Figure 4I**; sig. difference between conflict and no-conflict, p < 0.02; conflict, mean AUC = 0.98, 95% CI = [0.97,0.99]; no conflict, mean AUC = 0.94, 95% CI = [0.91,0.98]; representative population: conflict, mean AUC = 0.996, correctly classified 96.8% or 122/126 trials, no conflict AUC = 0.980, 87.3% correct or 55/63 trials, sig. change in correct classification likelihood, p < 0.04, 2 sample proportion test with pooled variance).

The increase in classification accuracy was due to an increase in the projection onto the correct response coding dimension (**Figure 4J-K**; p < 0.03, bootstrap test of the hypothesis that conflict minus no conflict is > 0; all trials: mean difference in projection onto task coding dimension = 0.24, 95% CI = [0.02, 0.48]; task 1 trials only: mean = 0.35, 95% CI = [-0.08, 0.79]; only task 2 trials: mean = 0.26, 95% CI = [-0.12, 0.63]; only task 3 trials: mean = 0.12, 95% CI = [-0.33, 0.54]). Thus, conflict increased the amount of correct response information in populations of neurons through shifting neural representations up the task coding axes, consistent with the amplification hypothesis. These population-level effects of conflict were qualitatively similar to what we observed in dlPFC (sig. difference in classification accuracy between conflict and no-conflict, p < 0.04, conflict, mean AUC = 0.97, 95% CI = [0.95,0.99]; no conflict, mean AUC = 0.92, 95% CI = [0.87,0.96]; increased projection onto correct response coding dimension during conflict, p < 0.2, mean difference in projection onto task coding dimension, all trial types = 0.30, 95% CI = [0.03, 0.58]).

### No abstract conflict coding axis in the population

Together, these results support the hypothesis that conflict amplifies neural coding of task variables within dACC. However, these results do not rule out the existence of a unified conflict axis. It thus remains possible that dACC both signal conflict *and* amplifies encoding of task variables. Therefore, we next asked whether there was a conflict detection axis in the population by examining the representational geometry of task variable and conflict coding dimensions in the region. Just as there are coding dimensions in the population that correspond to the task the subjects were performing, there are coding dimensions that correspond to the presence or absence of conflict during these tasks. In the amplification view, these must be at least partially aligned to the relevant task coding axis (**Figure 5A-B**). However, these conflict coding dimensions could also be at least partially aligned with each other. This would indicate that there is some average conflict coding vector that could be used to decode the presence or absence of conflict, regardless of the task. It would mean there was a conflict detection axis in the dACC population.

**Figure 5.**
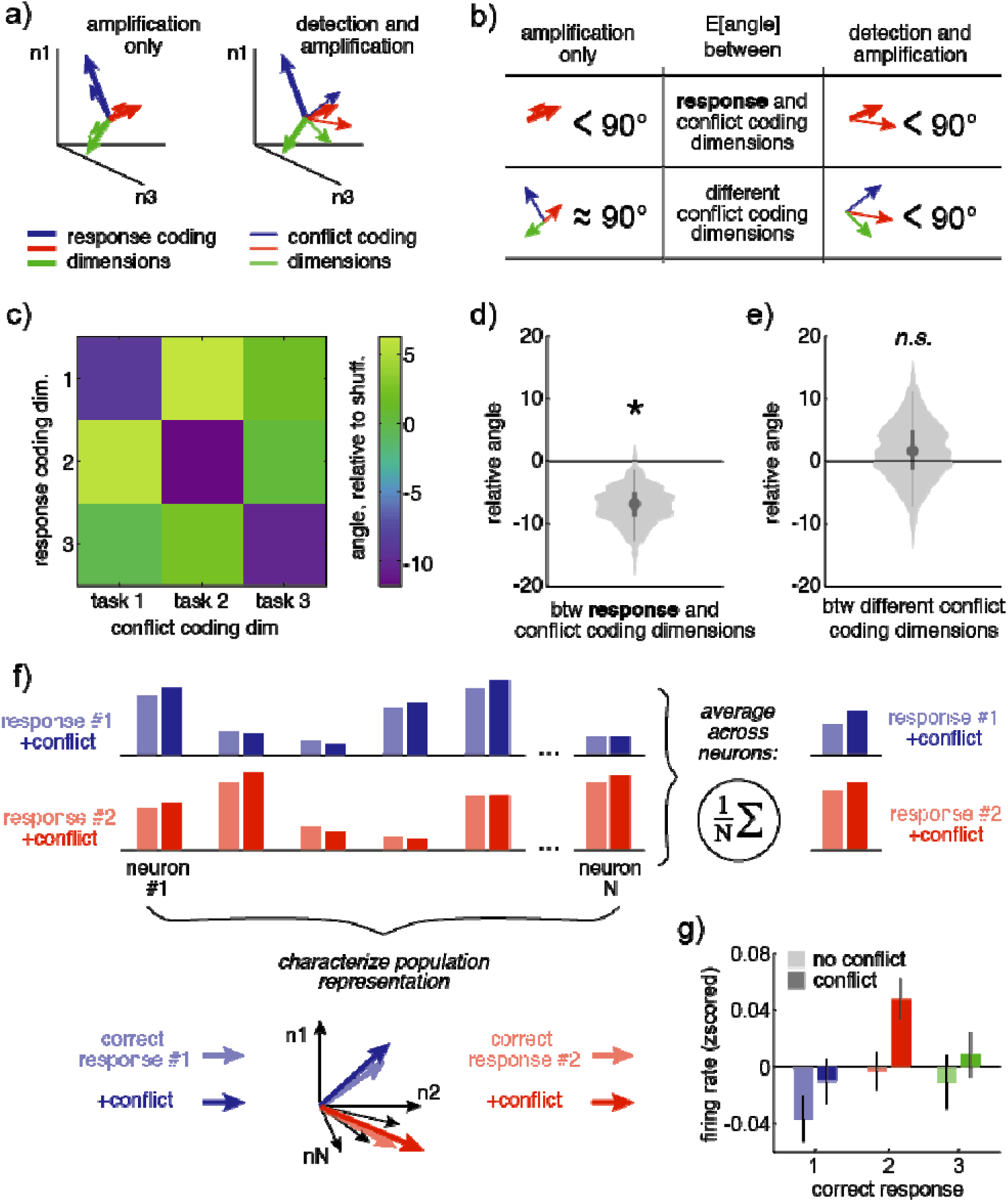
Relationship between task and conflict coding dimensions. A) A cartoon illustrating possible geometric relationships between the correct response coding dimensions and the population dimensions that encode the presence/absence of conflict during each task. Left) If conflict amplifies correct response information, conflict coding dimensions should be aligned with the matching correct response axes. Right) If dACC both explicitly detects conflict and amplifies correct responses, then there should be a shared conflict detection axis in the dACC population, which would mean that conflict coding axes will be at least partially aligned with each other. B) Predictions of the two hypotheses illustrated in A. For any amplification to occur, conflict coding dimensions must be somewhat aligned with the matching correct response coding dimensions. However, in the explicit conflict detection view, conflict coding dimensions would also be somewhat aligned with each other. C) The angle between each correct response coding dimension and conflict coding dimensions in the representative population. The diagonal structure indicates that conflict coding dimensions are aligned with matching response coding dimensions. Angles were normalized by subtracting the mean of label-permuted data, so 0 = no alignment. D) Distribution of angles between conflict coding dimensions and matched response coding dimensions across populations. E) Distribution of angles between the different conflict coding dimensions. F) A cartoon illustrating the central results. The population of neurons has a heterogenous pattern of activity for each correct response. Conflict modulates these patterns in different ways. Nevertheless, when averaging over neurons, conflict will generally increase activity, regardless of the correct response. However, one can also consider the whole pattern of activity across neurons, here illustrated as a neuron-dimensional vector. In this view, it becomes clear that the pattern of conflict modulation during one correct response is orthogonal to the pattern during another correct response. G) Conflict tended to increase average firing rates across neurons for each correct response condition, despite having orthogonal effects at the level of the pattern of population activity. Bars ± SEM across neurons.

We found that the conflict coding dimensions for each task were aligned with the task variable coding dimensions, both in the representative population (**Figure 5C**; more aligned than shuffled data, one-sided permutation test, p = 0.001) and across all the populations (**Figure 5D**; one-sided permutation test, p = 0.003). However, conflict coding dimensions were not aligned with each other (**Figure 5E**; not more aligned than shuffled data, one-sided permutation test, p > 0.5). Thus, while average neuronal firing rates tended to be higher when Ericksen conflict was present (**Figure 2A**) and this same trend was apparent regardless of the correct response (**Figure 5F-G**; mean difference, correct response 1 = 0.03 z-scored spikes/s, response 2 = 0.05, response 3 = 0.02), there was ultimately no explicit conflict detection axis in the dACC population. Instead, conflict amplified the encoding of the correct response.

## DISCUSSION

We sought to understand the neural basis of conflict processing by examining responses of neurons in human dACC and dlPFC collected in a conflict task. A previous paper from our team focused on spike-LFP relationships in this dataset and asked very different questions; the present one focuses on single unit activity (Smith et al., 2019). Here we show that the activity of dACC neurons tended to increase when conflict was present, consistent with most studies using mass action measures and with some recent neurophysiological studies (Sheth et al., 2012; Smith et al., 2019; Ebitz and Platt, 2015; Bryden et al., 2018; Michelet et al., 2015). Our major goal was to go beyond correlating neural activity with task variables, and to instead use *targeted dimensionality reduction* to determine what specific neuronal computations gave rise to this conflict signal. This method allowed us to directly compare and reject two major hypotheses in the literature, which we call the **explicit hypothesis** and the **epiphenomenal hypothesis** (Nakamura et al., 2005; Cole et al., 2009; Schall and Emeric, 2010; Mansouri et al., 2017; Cole et al., 2010; Kolling et al., 2016; Shenhav et al., 2016; Stuphorn et al., 2000). Instead, the data supported a third **amplification hypothesis**, that the effects of conflict are to amplify the encoding of task-relevant information across populations of neurons. Specifically, when conflict was present, the neural representation of the correct task-relevant sensorimotor responses was enhanced at the expense of irrelevant and incompatible responses (cf. Egner and Hirsch, 2005; Pastor-Bernier and Cisek, 2011; Cisek, 2007).

Attempts to determine the function of dACC have historically centered on identifying a specific executive role, that is, one that supports or modifies sensorimotor transformation but is external to and conceptually distinct from it (Paus, 2001; Bush et al., 2000; Ebitz & Hayden, 2016). This view is at least somewhat inconsistent with the growing literature identifying robust correlates of sensorimotor transformation in the region (e.g. Kennerley et al., 2011; Isomura et al., 2003; Johnston et al., 2007; Gemba et al., 1986; Hillman and Bilkey, 2010; Strait et al., 2016; Azab and Hayden, 2017; reviewed in Heilbronner and Hayden, 2016). That literature both suggests that dACC may have a sensorimotor role in addition to any executive role, and raises the broader question of how executive processes modulate sensorimotor transformations. It may helpful to think of dACC as one layer in a hierarchy of structures that can regulate goal-directed behavior by distributed changes in the gain of sensorimotor transformations (Cisek and Kalaska, 2010; Pezzulo and Cisek, 2016; Yoo and Hayden, 2018; Ebitz & Moore, 2017; Ebitz et al., 2019). Our results suggest that conflict is one of the executive processes that modulates sensorimotor encodings in this way. Note that conflict also has clear effects on the timing of action potentials, relative to ongoing local field potentials in this region, which may modulate the spiking effects we observed (Smith et al., 2019).

Amplification of task-relevant responses could push the system to focus on computations most relevant to the task at hand (Suzuki and Gottlieb, 2013; Finklestein et al., 2019; Ebitz et al., 2019; Ebitz et al., 2018; Egner & Hirsch, 2005). In this regard, we would draw an analogy between the effects of conflict we report in the prefrontal cortex and the effects of selective visual attention on sensory representations in the ventral visual stream (Desimone and Duncan, 1995; Desimone, 1996; McAdams and Maunsell, 1999). Attention is the enhanced representation of behaviorally-relevant stimuli at the expense of other stimuli. Representations of stimuli naturally compete for control of behavior, and attention functions to bias this competition towards behaviorally relevant representations. Notably, the competition between representations does not stop at the rostral pole of the temporal lobe, but continues through to the motor system (Cisek and Kalaska, 2010; Cisek, 2007; Cisek, 2012). While it is not clear whether the same computations are involved in biasing competition between sensory representations in extrastriate cortex, motor representations in motor cortices, or sensorimotor representations in association cortices, it is clear that there could be a continued benefit in a biasing process that can tip the scales towards favored action at any point throughout the sensorimotor transformation. Further, visual attention may produce shifts in population-level stimulus representations in extrastriate cortex that resemble the shifts that conflict produces in sensorimotor representations in the prefrontal cortex (Cohen and Maunsell, 2010). Thus, it seems prudent to consider the possibility that cognitive processes like conflict may invoke computational processes that resemble those that bias competition between sensory representations in extrastriate cortex (Cisek, 2007; Michelet et al., 2010; Pastor-Bernier and Cisek, 2011; Yoo et al., 2018). In other words, our findings are consistent with the idea that the brain uses conserved computational processes to solve similar problems in different ends of the brain (Yoo and Hayden, 2018; Hunt et al., 2017).

Our results highlight the differences between dACC and dlPFC. These two regions are strongly interconnected, and are both strongly implicated in many executive functions. This relatedness does not necessarily imply that they have identical roles, however (Smith et al., 2019; Hunt et al., 2018). Indeed, anatomy and functional studies both motivate the hypothesis that these regions may have a hierarchical relationship (Shenhav et al., 2017; Miller and Cohen, 2000; MacDonald et al., 2000) as do at least some physiological studies (Hunt et al., 2018). In this hierarchical view, the increase in conflict modulation that we observed in dACC neurons may occur because the region responds to conflict at an earlier and more abstract level of the hierarchy, while dlPFC is less modulated by conflict because it is later and presumably more effector-specific. Of course, the hierarchical view does not require that regions must have strict functional differences, but instead a gradual shift in function along a hierarchy that transforms sensations to actions (Hunt et al., 2017; Yoo and Hayden, 2018).

The differences between the Eriksen/flanker and Simon/response conflict effects we report here echo earlier findings from human EEG (Van Veen et al., 2001) and primate neurophysiology (Ebitz and Platt, 2015). These earlier studies report that that conflict encoding can differ depending on whether the conflict is between responses / stimuli (Van Veen et al., 2001) or between responses / task sets (Ebitz and Platt, 2015). The two forms of conflict in our task have some intuitive similarities to the distinction between the different forms of conflict in these previous studies. However, the overlap is unlikely to be perfect - as Van Veen et al., showed, the flanker task can elicit both stimulus and response conflict depending on the condition. Nonetheless, this study supports the conclusions drawn by these previous studies—that different types of conflict may not have unitary effects on brain activity.

Our results do not answer the important question of where the cognitive control for response to conflict originally comes from. We see two possibilities, both consistent with our data. First, there may be another region – distal to dACC – that detects conflict and controls responses of dACC task-selective neurons. Second, there may be no single region that functions as a central executive. Certainly, it is possible to build executive systems that lack a central controller (Eisenreich et al., 2017). For example, ant colonies – a canonical distributed decision-making system – show what may be described as executive control, even in the absence of a central executive (Franks et al., 2002; Franks et al., 2003). Future work, including modeling, will be needed to disambiguate these two hypotheses.

## METHODS

### Subjects and ethics statement

We studied two cohorts of subjects. Cohort 1 consisted of 7 patients (1 female) with medically refractory epilepsy who were undergoing intracranial monitoring to identify seizure onset regions. Before the start of the study, these subjects were implanted with stereo-encephalography (sEEG) depth electrodes using standard stereotactic techniques. One or more of the sEEG electrodes in this cohort spanned dorsolateral prefrontal cortex (dlPFC) to dorsal anterior cingulate cortex (dACC; Brodmann’s areas 24a/b/c and 32), providing LFP recordings from both regions, as well as single unit recordings in dACC (see below; Data Acquisition).

Cohort 2 consisted of 9 patients: 8 (2 female) with movement disorders (Parkinson’s disease or essential tremor) who were undergoing deep brain stimulation (DBS) surgery, and one male patient with epilepsy undergoing intracranial seizure monitoring. The entry point for the trajectory of the DBS electrode is typically in the inferior portion of the superior frontal gyrus or superior portion of the middle frontal gyrus, within 2 cm of the coronal suture. This area corresponds to dlPFC (Brodmann’s areas 9 and 46). The single epilepsy patient in this cohort underwent a craniotomy for placement of subdural grid/strip electrodes in a prefrontal area including dlPFC.

All decisions regarding sEEG and DBS trajectories and craniotomy location were made solely based on clinical criteria. The Columbia University Medical Center Institutional Review Board approved these experiments, and all subjects provided informed consent prior to participating in the study.

### Behavioral Task

All subjects performed the multi-source interference task (MSIT; **Figure 1A**). In this task, each trial began with a 500-millisecond fixation period. This was followed by a cue indicating the ***correct response*** as well as the ***distractor response***. The cue consisted of three integers drawn from {0, 1, 2, 3}. One of these three numbers (the “***correct response cue***”) was different from the other two numbers (the “***distractor response cues***”). Subjects were instructed to indicate the identity of the correct response number on a 3-button pad. The three buttons on this pad corresponded to the numbers 1 (left button), 2 (middle) and 3 (right), respectively.

The MSIT task therefore presented two types of conflict. Simon (motor spatial) conflict occurred if the correct response cue was located in a different position in the cue than the corresponding position on the 3-button pad (e.g. ‘0 0 1’; target in right position, but left button is correct choice). Eriksen (flanker) conflict occurred if the distractor numbers were possible button choices (e.g. ‘3 2 3’, in which “3” corresponds to a possible button choice; vs. ‘0 2 0’, in which “0” does not correspond to a possible button choice).

After each subject registered his or her response, the cue disappeared and feedback appeared. The feedback consisted of the target number, but it appeared in a different color. The duration of the feedback was variable (300 to 800 milliseconds, drawn from a uniform distribution therein). The inter-trial interval varied uniformly randomly between 1 and 1.5 seconds.

The task was presented on a computer monitor controlled by the Psychophysics Matlab Toolbox (www.psychtoolbox.org; The MathWorks, Inc). This software interfaced with data acquisition cards (National Instruments,) that allowed for synchronization of behavioral events and neural data with sub-millisecond precision.

### Data Acquisition and preprocessing

Single unit activity (SUA) was recorded from microelectrodes using 3 different techniques. In Cohort 1, the dlPFC-dACC sEEG electrodes were Behnke-Fried macro-micro electrodes (AdTech Medical). These electrodes consist of a standard clinical depth macroelectrode shaft with a bundle of eight shielded microwires that protrude ∼4 mm from the tip (IRB-AAAB6324). These 8 microwires are referenced to a ninth unshielded microwire.

Cohort 2 provided dlPFC SUA, although it used a combination of two techniques. The DBS surgeries were performed according to standard clinical procedure, using clinical microelectrode recording (Frederick Haer Corp.). Prior to inserting the guide tubes for the clinical recordings, we placed the microelectrodes in the cortex under direct vision to record from dlPFC, (IRB-AAAK2104). The epilepsy implant in Cohort 2 included a Utah-style microelectrode array (UMA) implanted in dlPFC (IRB-AAAB6324). In all cases, data were amplified, high-pass filtered, and digitized at 30 kilosamples per second on a neural signal processor (Blackrock Microsystems, LLC).

SUA data were re-thresholded offline at negative four times the root mean square of the 250 Hz high-pass filtered signal. Well-isolated action potential waveforms were then segregated in a semi-supervised manner using the T-distribution expectation-maximization method on a feature space comprised of the first three principal components using Offline Sorter (OLS) software (Plexon Inc, Dallas, TX; USA). The times of threshold crossing for identified single units were retained for further analysis.

#### Additive effects of Simon and Ericksen conflict

We determined what effect the combination of Ericksen and Simon conflict had on dACC activity by comparing the fit of the following three generalized linear models. First, we considered a 4-parameter “full model”, which independently measured the contribution of Ericksen conflict (C^E^; coded as 1 when the correct response and Ericksen distractor response were in conflict, 0 otherwise), Simon conflict (C^S^), and the combination of both (C^B^; coded as 1 if and only if C^E^ and C^S^ were both true). This model 1) made no assumptions about the relative effects of Ericksen and Simon conflict and 2) also allowed for superadditive or subadditive effects when both forms of conflict co-occurred.

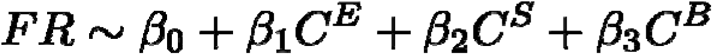

For the second model, the “independent model”, we dropped the sub-/super-additive term, leaving a simplified, 3-parameter model. This model would be a sufficient explanation for the data if the dACC response to the combination of Ericksen and Simon conflict was simply the sum of the two types of conflict independently.

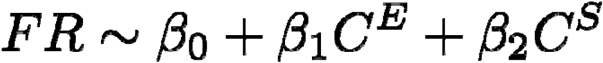

Finally, in the third, “additive model”, we dropped the term that allowed Simon and Ericksen conflict to have different effects (i.e. we assumed that *β*_1_ = *β*_2_in our previous model), leaving a 2-parameter model. This model would be a sufficient explanation for the data if Ericksen and Simon conflict have both identical and additive effects on the dACC population.

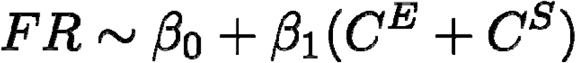

We used standard model comparison (Burnham & Andersen, 2010) to determine whether each simplifying assumption could be made with no loss of information. Models were fit to z-scored firing rates that were condition-averaged within neurons (9 data points per neuron, reflecting all combinations of the 3 correct response, 3 Ericksen distractors, and 3 Simon distractors) and offset terms were included for each neuron (number of neurons-1 offset terms), though the z-scoring ensured that the results did not depend on including cell identity terms.

**Table S1.**
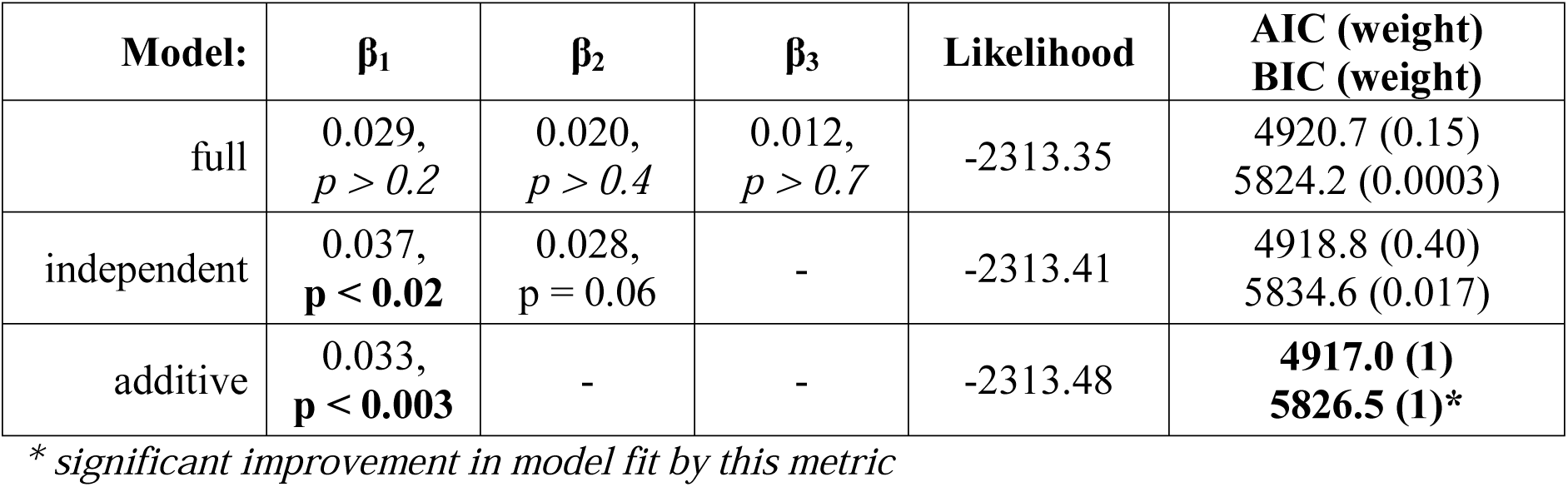
Additive effects of Simon and Ericksen conflict.

#### Task, distractor, and conflict tuning

To determine how frequently correct response, distractor response, and conflict tuning co-occurred within individual cells, we used the following ANOVA:

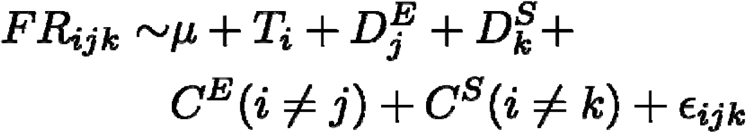

Where FR_ijk_ is the average firing rate of the cell for the ith correct response, with the jth Ericksen distractor response, and kth Simon distractor response. Here, the T term models the effect of correct responses on neural activity, the D^E^ and D^S^ terms model, respectively, the effects of Ericksen and Simon distractor responses, and the C^E^ and C^S^ terms model the effects of conflict, meaning the mismatch between correct and distractor responses for Ericksen and Simon distractors, respectively.

#### Residuals

In the epiphenomenal hypothesis, conflict signals are an emergent consequence of co-activating pools of neurons that are tuned for different responses (**Figure 4A**). This implies that we should be able to predict activity in conflict conditions from the activity under different task and distractor conditions. Systematic deviations from these predicted values would indicate some pattern that could not have emerged because of summed activity due to task and distractor activations without some form of systematic nonlinearity (which we have no reason to expect a priori in dACC). Within each neuron, we calculated the expected firing rate for each task condition, marginalizing over distractors, and for each distractor, marginalizing over tasks. Then we estimated the expected activity for each combination of task and distractor by summing these estimates for task and distractor (**Figure 3C**). Subtracting this expectation from the observed pattern of activity left the residual activity that could not be explained by the linear co-activation of task and distractor conditions.

#### Pseudopopulations

To estimate how conflict affected neuronal population activity, we generated pseudopopulations by combining the activity of neurons that were recorded largely separately (Churchland et al., 2012; Machens et al., 2010; Mante et al., 2013; Meyers et al., 2008; Ebitz et al., 2019; Sleezer et al., 2016). Within each task condition (combination of correct response and distractor response), firing rates from separately recorded neurons were randomly drawn with replacement to create a pseudotrial firing rate vector for that task condition, with each entry corresponding to the activity of one neuron in that condition. Pseudotrial vectors were then stacked into the trials-by-neurons pseudopopulation matrix. Nineteen pseudotrials were drawn for each condition, based on the observation that a minimum of 75% of conditions had at least this number of observations, though results were identical for different choices of this value (± 5 trials). All effects were confirmed via bootstrap tests across 1000 randomly re-seeded pseudopopulations. In addition, some analyses are reported with a “representative” pseudopopulation. This was the pseudopopulation that most closely matched the average condition means across 1000 random samples (i.e. the pseudopopulation seed that minimized root mean squared error from the vector of condition projection averages). These analyses focus on Eriksen conflict because this form of conflict had the larger effect on response time and caused a significant increase in the average activity of dACC neurons. Similar results were obtained for Simon conflict (data not shown).

#### Targeted dimensionality reduction

To determine how conflict affected population activity along task-coding dimensions, we used a form of targeted dimensionality reduction based on multinomial logistic regression (Ebitz, et al., 2018; Peixoto et al., 2018). Targeted dimensionality reduction is a class of methods for re-representing high-dimensional neural activity in a small number of dimensions that correspond to variables of interest in the data (Cohen and Maunsell, 2010; Cunningham and Yu, 2014; Mante et al., 2013; Peixoto et al., 2018; Ebitz et al., 2019). Thus, unlike principle component analysis—which reduces the dimensionality of neural activity by projecting it onto the axes that capture the most variability in the data—targeted dimensionality reduction reduces dimensionality projecting activity onto axes that encode task information or predict behavior.

Here, we were interested in how conflict changed task coding, so we first identified the axes in neural activity discriminated the three correct responses. We used multinomial logistic regression to find the separating hyperplanes in neuron-dimensional space that best separated the neural activity for one correct response (i.e. button press 1) from activity during the other correct responses (i.e. not button press 1). Formally, we fit a system of three logistic classifiers:

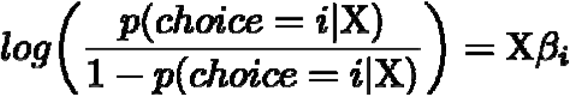

Where X is the trials by neurons pseudopopulation matrix of firing rates and βi is the vector of coefficients that best differentiated neural activity on trials in which a choice matching category i is chosen from activity on other trials (fit via regularized maximum likelihood). The separating hyperplane for each choice i is the vector (a) that satisfies:

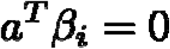

Meaning that βi is a vector orthogonal to the separating hyperplane in neuron-dimensional space, along which position is proportional to the log odds of that correct response: this is the the coding dimension for that correct response. By projecting a pseudotrial vector x onto this coding dimension, we are essentially re-representing the trial in terms of its distance from the separating hyperplane corresponding to task i. Projecting that trial onto all three classifiers, then re-represents that high-dimensional pseudotrial in three dimensions—each one of which corresponds to the coding dimension of a different response.

We used an identical approach to identify the coding dimensions corresponding to each distractor. To identify coding dimensions corresponding to task conditions (combination of 3 correct responses and 3 distractor responses, if present), we used the same approach to classify the 12 task conditions.

## Acknowledgements

This work was supported by NIH R01 MH106700, NIH K12 NS080223, NIH S10 OD018211, NIH R01 NS084142, NIH R01 DA038615, the Brain and Behavior Foundation, and the Dana Foundation. Special thanks to Camilla Casadei, David K. Peprah, Yagna Pathak, and Timothy G. Dyster for coordination and data collection efforts. The funders had no role in study design, data collection and analysis, decision to publish, or preparation of the manuscript.

